# Human deleterious mutation rate slows adaptation and implies high fitness variance

**DOI:** 10.1101/2023.09.01.555871

**Authors:** Joseph Matheson, Ulises Hernández, Jason Bertram, Joanna Masel

## Abstract

Each new human has an expected *U*_*d*_ = 2-10 new deleterious mutations. Using a novel approach to capture complex linkage disequilibria from high *U*_*d*_ using genome-wide simulations, we confirm that fitness decline due to the fixation of many slightly deleterious mutations can be compensated by rarer beneficial mutations of larger effect. The evolution of increased genome size and complexity have previously been attributed to a similarly asymmetric pattern of fixations, but we propose that the cause might be high *U*_*d*_ rather than the small population size posited as causal by drift barrier theory. High within-population variance in relative fitness is an inevitable consequence of high *U*_*d*_∼2-10 combined with inferred human deleterious effect sizes; two individuals will typically differ in fitness by 15-40%. The need to compensate for the deluge of deleterious mutations slows net adaptation (i.e. to the external environment) by ∼13%-55%. The rate of beneficial fixations is more sensitive to changes in the mutation rate than the rate of deleterious fixations is. As a surprising consequence of this, an increase (e.g. 10%) in overall mutation rate leads to faster adaptation; this puts to rest dysgenic fears about increasing mutation rates due to rising paternal age.

## Introduction

The average human begins life with around a hundred new mutations not found in their parents, and geneticists have long worried about the effects of the resulting “mutation load” on human health ^1–6^. Lesecque et al. ^7^ assumed that mutations are deleterious only in the 55% of the 6 × 10^9^ diploid genome that is not dominated by inactive transposable elements, which evolves 5.7% slower than the rest of the genome due to this constraint, at a point mutation rate of 1.1 × 10^−8^; this yields an estimated rate of deleterious mutations of 0.55 × 6 × 10^9^ × 0.057 × 1.1 × 10^−8^ = 2.1 per replication. This estimate is conservative: some mutations to the other 45% of the genome are deleterious, more recent estimates of the human point mutation rate have 1.1 × 10^−8^ as the lower bound of a 95% confidence interval ^8^, and non-point mutations and beneficial mutations are neglected. Some therefore argue that the deleterious mutation rate is as high as ten ^9,10^. Mutation rates of this order are not unique to humans ^8,11^.

Three fears about the effects of high deleterious mutation rates have haunted human genetics, namely that: 1) we do not understand how population persistence is possible ^12^; 2) the health burden caused by this deluge of deleterious mutations is not shared equally by all humans; and 3) recent increases in mutation rate and/or decreases in natural selection will lower population fitness at mutation-selection balance, with potentially catastrophic consequences for health systems ^1–3,6^. The second fear has a dark legacy of attempting to identify more vs. less 'burdened' humans ^13,14^.

Most substantial theoretical treatment has focused on the first fear. Some unconditionally deleterious mutations of smaller effect size will inevitably fix ^15,16^, producing cumulative degradation termed “Ohta’s ratchet” ^17^. But rarer, larger-effect beneficial fixations can compensate, enabling population persistence ^18– 20^. Given a distribution of fitness effects in an infinite sites model, fixed deleterious mutations tend to have smaller *s* ^21^ while beneficial fixations have larger *s* ^22^. Asymmetry between deleterious and beneficial fixations matches known features of molecular adaptation. E.g., many proteins accumulate small deleterious mutations that slightly inhibit folding, which can be compensated for by a novel or improved or overexpressed chaperone protein ^23^. Similarly, many poorly splicing introns can be ameliorated by the evolution of a better spliceosome ^24^. An asymmetric pattern of adaptation has been inferred in influenza ^25^.

While the second fear has received little rhetorical attention from population geneticists, it has received incidental theoretical treatment as “load” in the context of the first fear. Galeota-Sprung *et al*. ^26^ found that the expected variance in fitness is 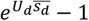 in the absence of linkage disequilibrium or epistasis. They conclude that, unlike load metrics that compare to an ideal individual, variance in fitness is modest, and not a problem for population persistence. They did not discuss the societal implications of variance in fitness. They underestimated variance by using unrealistically low values of 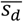 (see Results), and by neglecting linkage disequilibrium that makes purging harder. Similarly, warnings from prominent geneticists of a coming health crisis use one-locus models of mutation-selection balance ^1–3,6^, raising similar questions of the role of linkage disequilibrium; they also neglect large-effect beneficial mutations.

Quantitative treatment of the three fears requires incorporating both beneficial mutations, and emergent linkage disequilibria ^27^. Linkage disequilibria become especially pronounced when *U*_*d*_ > 1 ^28,29^. Unfortunately, simulating this is computationally challenging. One remedy of convenience when simulating *U*_*d*_ > 1 is to neglect beneficials, and instead periodically re-normalize relative fitness to cosmetically remove ongoing degradation (e.g. compare Fig. S2 to Fig. 2 in ^30^). Alternatively, most previous studies of mutation load used unrealistically low mutation rates (*U*_*d*_ < 1), either directly ^31–37^, or by assuming independent loci ^30,38–40^.

Simulation is essential to capture the full complexities of multilocus linkage disequilibrium. Most forward time simulation methods hold a product such as *sN* constant by rescaling *N* to be smaller and *s* to be larger, speeding computation ^41^. But this reduces the number of segregating mutations *UNτ* (where *τ* is the expected sojourn time), which understates the impact of linkage disequilibria. We simulate a census population size of 20,000, producing a human-like level of neutral diversity (*N*_*e*_ ∼7500), with emergent linkage disequilibrium. To avoid computationally costly individual tracking of the enormous number of segregating sites, we track only the combined fitness effects within ‘linkage blocks’ between recombination hotspots, using a novel algorithm. Our approach allows us to recover information about fixed mutations via tree-sequence recording ^42^, albeit at a cost to runtime.

We use simulations to quantify, given realistic linkage disequilibria, the fluxes of beneficial and deleterious fixations, the speed of adaptation, and variation in fitness within populations with different mutation rates. This allows us to quantitatively treat all three load-based fears.

## Methods

Each individual has two characteristics: a genome, and a fitness value derived from it. Each individual’s genome is represented as two haplotypes, each an array of *L* non-recombining ‘linkage blocks’, divided into 23 chromosomes. Each linkage block consists of a floating-point variable *l*_*j*_, which summarizes the fitness effects of all mutations that occurred in the history of that linkage block, such that *l*_*j*_ = *∏*_*i*_(1 + *s*_*i*_). We assume a multiplicative form of co-dominance and no epistasis, such that *w*_*i*_ = 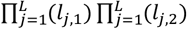 where *l*_*j*,1_ and *l*_*j*,2_ refer to the effects of linkage block *j* in haplotypes 1 and 2, respectively. Note that this computationally convenient choice is not precisely equivalent to a typical codominance model, where 1 + *s*_*i*_ is the fitness of a homozygote and 1 + *s*_*i*_*h*_*i*_ is the fitness of a heterozygote. While co-dominance is unrealistic for strongly deleterious mutations, which are often highly recessive, it is reasonable for the small-effect deleterious mutations that drive Ohta’s ratchet ^43–45^.

In addition to independent assortment of chromosomes, recombination occurs at hotspots between linkage blocks via crossing-over events between homologous chromosomes. We simulate exactly two recombination events per chromosome per meiosis, matching data for humans ^46^, although we don’t explicitly simulate a centrosome. Representing a genome as a set of ‘linkage blocks’ is a good approximation of population genetics in non-microbial species ^47–49^. Realistic values of *L* in humans are in the range of 10^5^-10^6 50–54^. Once *L* ≥ 50 × 23 = 1150, results converge (Supplementary Figure 1), so for computational efficiency we use *L* = 50 × 23. This simplification should overestimate the effect of linkage between selected mutations, which is conservative with respect to the ability of beneficial mutations to counteract load.

Following recombination, we sample the number of new deleterious mutations in the gamete from a Poisson distribution with mean *U*_*d*_. Our distribution of fitness effects is based on a large empirical study of Europeans ^55^, who fitted a gamma distribution for 2*N*_*e*_ *sh* with mean −224.33, shape parameter α = 0.169 and scale parameter *β* = 1327.4. After drawing a value of 2*N*_*e*_*sh* from this distribution, we rescale to *sh* using their inferred *N*_*e*_ = 11,823. We use the *sh* value drawn from this distribution as our *s*_*d*_ value; its mean is 0.009. We sample the number of new beneficial mutations from a Poisson distribution with mean *U*_*b*_, and fitness effects drawn from an exponential distribution with mean *s*_*b*_ (again, this is the fitness effect in the heterozygote). We explore a range of values for *U*_*b*_ and *s*_*b*_ that we consider *a priori* plausible: *U*_*b*_ ∼ 0.0001-0.01 and *s*_*b*_ ∼ 0.001-0.01.

We simulate a Moran model with constant population size *N*. An individual chosen uniformly at random dies each time step and is replaced by a child produced by two parents, who are chosen with probability proportional to their fitness *w*_*i*_. Each generation consists of *N* time steps. The fitnesses of the population are stored in an unsorted array — in a naïve implementation, exchanging an element to represent a birth and death would be rapid, but sampling proportional to fitness would be *O*(*N*). The current fastest forward-time genetic simulation tools for large population sizes (e.g. fwdpy ^56,57^ and SLiM ^58^) preprocess cumulants each generation in a Wright-Fisher model; this speeds up sampling from the fitness array, and while the processing algorithm is *O*(*N*), it only needs to be performed once per generation. We instead use a binary indexed tree ^59^ to sample fitnesses efficiently according to the cumulative probability distribution — both updating and sampling from the tree are *O*(*log N*). Our scheme is expected to have similar efficiency but is intended to be useful for future expansions of this approach to absolute fitness and more complex life history models ^60,61^, e.g. to allow better treatment of reproductive compensation ^62^.

We initialize the population with mutationless individuals, then conduct a ‘burn-in’ phase during which variation increases to stable levels (Supplementary Figure 2). We end the burn-in phase 500 generations after a linear regression of the variance in fitness over the last 200 generations produces a slope less than an arbitrarily chosen low value of 0.007/*N* that we visually confirmed to perform well (e.g. Supplementary Figure 2). The length of the burn-in phase does not strongly depend on *N* (Supplementary Figure 3).

We calculate the net fitness flux from each simulation as the slope of the regression of log mean population fitness on time after burn-in (Supplementary Figure 2, black slope following dashed line). To numerically solve for a specified net fitness flux for Figure 1, we varied *s*_*b*_ while holding *U*_*b*_ constant. Our algorithm finds values of *s*_*b*_ that bracket the target net fitness flux, then uses a bisection method until it finds a value of *s*_*b*_ that is within ±0.00005 of the target. In practice, there was little stochasticity in the regression slope (which averages out stochasticity in the timecourse), and so this relatively deterministic method performed well.

**Figure 1.**
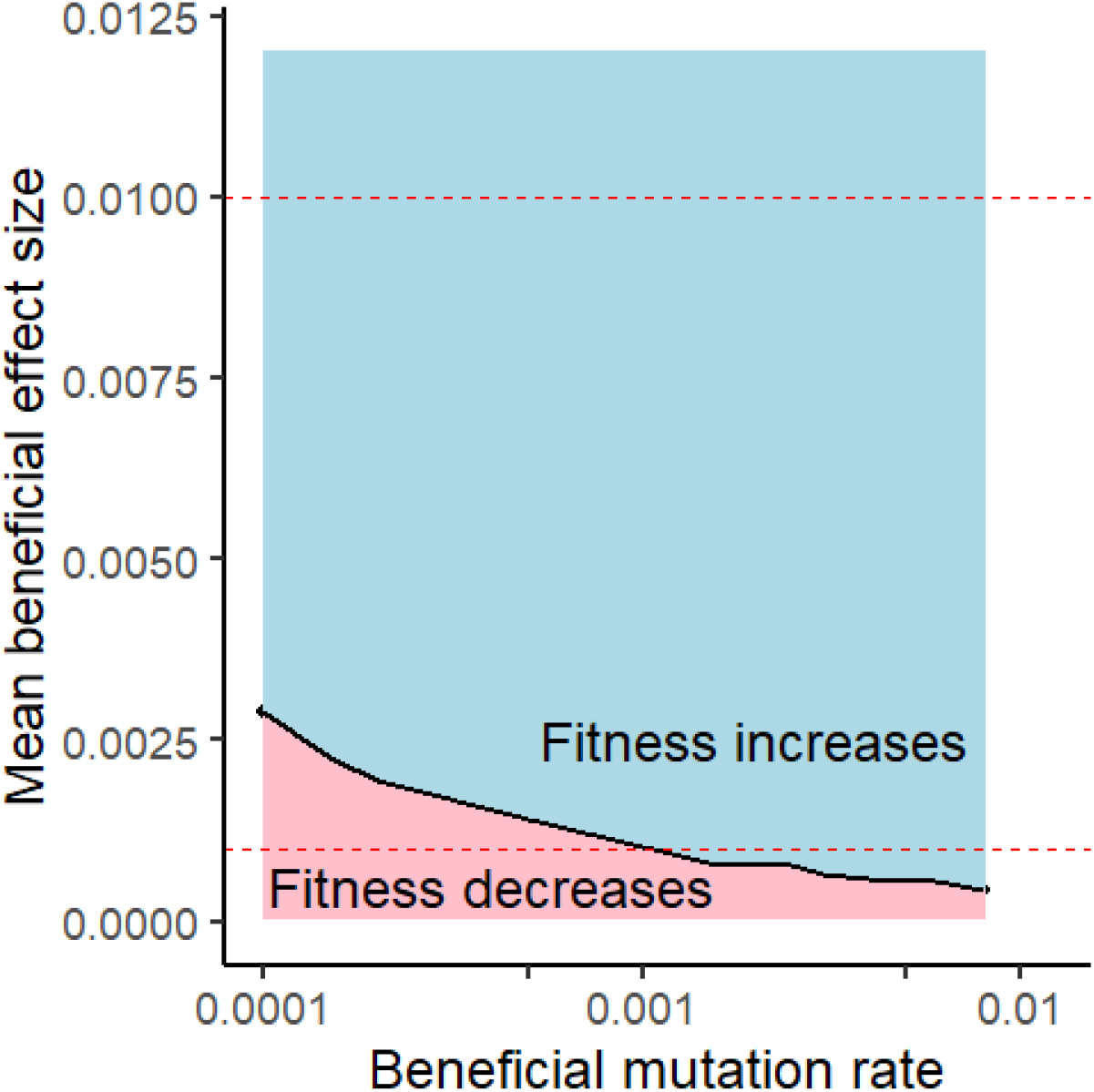
Relatively rare and mild beneficial mutations are sufficient to counteract a deluge of slightly deleterious mutations accumulating under Ohta’s ratchet. Black line shows combinations of beneficial mutation parameters that produce zero net fitness flux. All populations simulated with *N* = 20,000, *U*_*d*_ = 2, and 23 chromosomes with 50 linkage blocks per chromosome. Red dashed lines show plausible upper and lower estimates of the mean effect size of new beneficial mutations in humans that we deemed *a priori* plausible; almost all of the resulting parameter value range produces positive net fitness flux. The range for *U*_*b*_ was chosen to be broad, including values that are likely too low.

Although the census population size *N* is a parameter of our model, the effective population size *N*_*e*_ is not, but rather emerges over the course of a given simulation. To estimate it, we recorded tree-sequences using the tskit package ^42^, then used msprime ^63^ to retroactively add neutral mutations after each simulation. We did this only for one parameter combination involving realistically high *N*, due to the significant computational cost of this procedure; this was 23 chromosomes, 50 linkage blocks per chromosome, *N* = 20,000, *U*_*d*_ = 2, *U*_*b*_ = 0.002, and *s*_*b*_ = 0.0025. These parameter values produce only a small excess of adaptation above that needed to counter Ohta’s ratchet (Figure 1). We calculate *N*_*e*_ using neutral heterozygosity under an infinite-alleles model. The choice of neutral mutation rate will not affect estimated *N*_*e*_, so long as it is small enough to avoid multiple mutations at the same site; we arbitrarily chose 1.0 × 10^−6^ per linkage block, or 1.15 × 10^−4^ per haploid genome. This produced *N*_*e*_ ∼7500, on the order of effective population sizes inferred for ancestral human populations ^64^. For comparison, similar simulations with *U*_*b*_ = 0 (i.e. with background selection alone and declining relative fitness), produce *N*_*e*_∼16,000 ^28^. This is consistent with other results, using the simulation approach developed here, showing that *N*_*e*_ is primarily lowered by interactions between beneficial and deleterious mutations ^29^.

Tree sequence recording also tracks all non-neutral mutations, so that we can identify those that fixed and thus determine the degree of asymmetry in the effect sizes of fixed mutations. Note that without tree-sequence recording, this information would be inaccessible due to the way we summarize the fitness of many mutations within linkage blocks. However, using tree-sequence recording for all non-neutral mutations significantly increases the computation time of simulations. For a representative case with *N* = 20,000, *U*_*b*_ = 0.001, *U*_*d*_ = 2, *s*_*b*_ = 0.001, 23 chromosomes with 50 linkage blocks each, and all other details as described above, a single simulation of 200,000 generations took 12 hours, 45 minutes without tree-sequence recording and 58 hours, 42 minutes with tree-sequence recording. When we are solving for the parameters that produce a target value of net fitness flux, we therefore do not use tree sequence recording.

## Results

The mean beneficial effect size 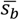 is more important than the beneficial mutation rate *U*_*b*_ for achieving net positive fitness flux (relatively horizontal contour of zero net flux in Figure 1, despite log scale on x-axis). While there is great uncertainty in the true values of *U*_*b*_ and 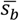, the 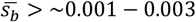 range required for population persistence seems entirely plausible, demonstrating that modest assumptions about beneficial mutations are sufficient to rescue populations from Ohta’s ratchet.

Beneficial fixations are of much greater magnitude than deleterious fixations. In the representative case shown in Figure 2, fixations have 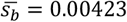 (more than four times as large as *de novo* beneficial mutations) 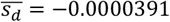 (almost three orders of magnitude smaller than the mean of −0.009 for *de novo* deleterious mutations). 56,255 deleterious and 6,387 beneficial fixations occurred over the 200,000 generations simulated for Figure 2.

**Figure 2.**
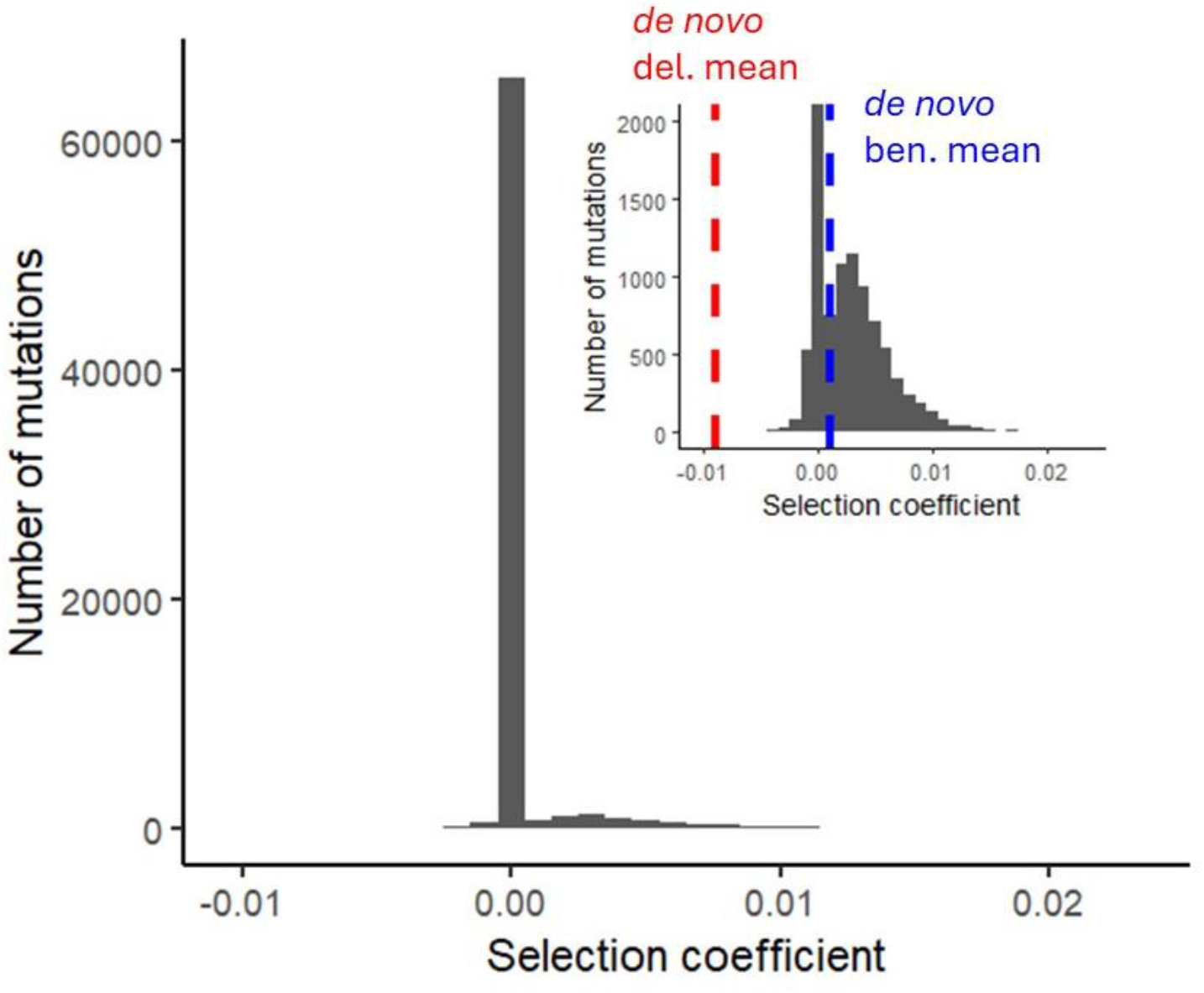
Damage from small effect deleterious fixations is balanced by far fewer but larger effect beneficial fixations. The distribution of effect sizes of fixed mutations is shown in bins of 0.001 at the end of 200,000 generations, in a population with *N* = 20,000, *U*_*b*_ = 0.001, *U*_*d*_ = 2, *s*_*b*_ = 0.001, and 23 chromosomes with 50 linkage blocks each. Inset truncates the y-axis to see detail outside the mode of slightly deleterious fixations. Effect sizes are per allele copy.

To adapt to a changing environment, a population must generate positive net fitness flux, i.e. beneficial fixations above and beyond those required to counterbalance Ohta’s ratchet. Rapid adaptation is shown in purple in Figure 3, slow adaptation in green, and an inability even to counter deleterious fixations in black. While resistance to mutational degradation remains reasonably robust to high *U*_*d*_ (few black squares), the need to compensate for Ohta’s ratchet drives down the rate of adaptation to a changing external environment by an average of ∼13%, ∼35%, and ∼55% for *U*_*d*_ of 2, 5, and 10, respectively, relative to *U*_*d*_ = 0.

**Figure 3.**
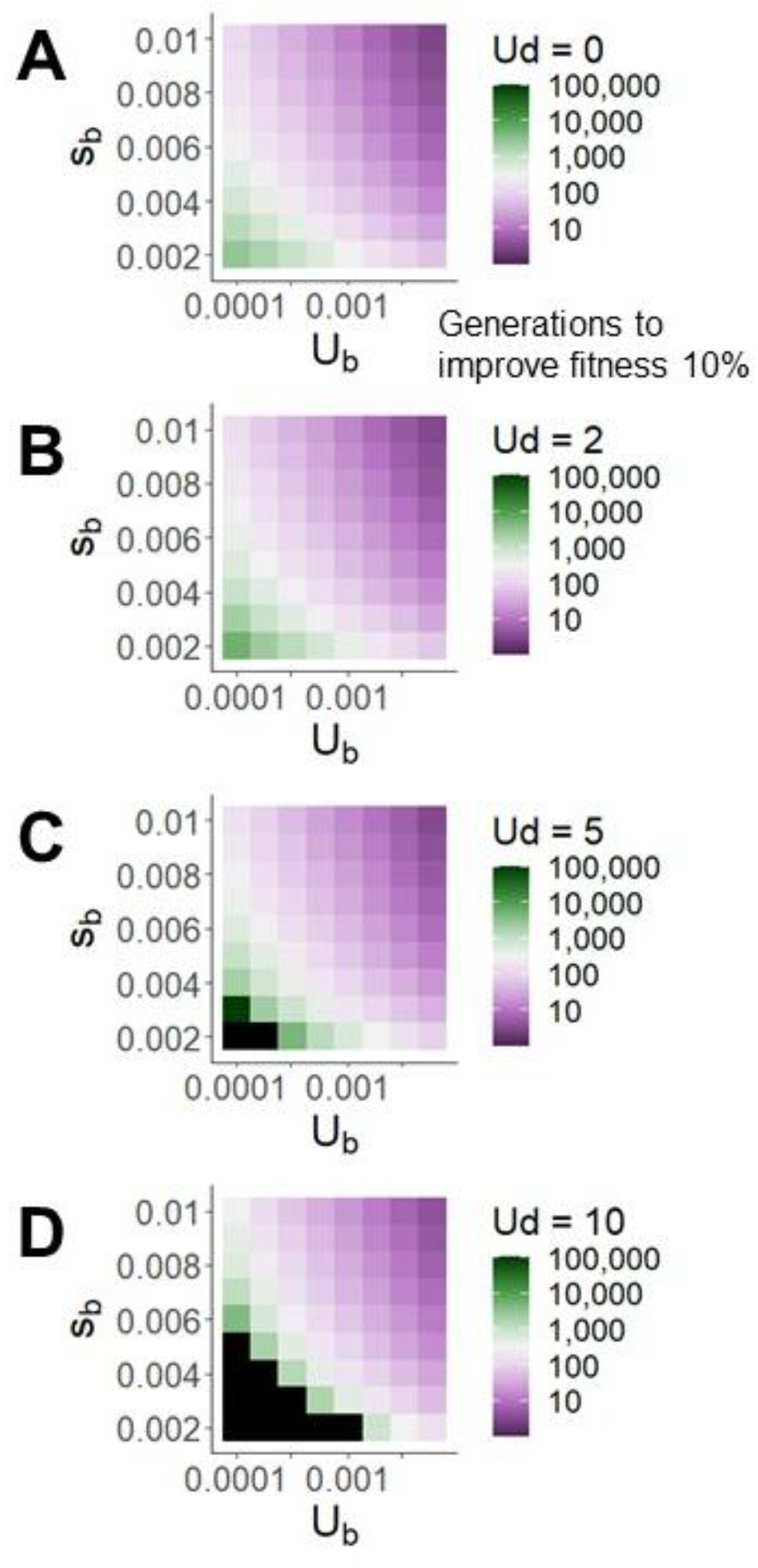
Deleterious mutations appreciably but modestly slow adaptation, visualized as the number of generations required for population mean fitness to increase by 10%. Black boxes indicate simulations with net fitness flux < 0.

Population geneticists have raised concerns about the increase in the human mutation rate ^1^, in particular due to increased age at paternity ^2^. Mean paternal age in the U.S. increased from 27.4 to 30.9 years between 1972 and 2015 ^65^, which would add about 7 *de novo* mutations to the 70-100 experienced by typical human gametes, or a 7-10% increase in the mutation rate ^66^. We simulated a 10% increase in the mutation rate, for both deleterious and beneficial mutations, for a reference population with *U*_*b*_ = 0.002, 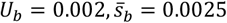, and other parameters, including baseline *U*_*d*_ = 2, as in Figure 1. Surprisingly, populations with increased mutation rates took only 127 generations to increase their fitness by 10%, compared to 151 generations for the baseline population. In other words, because beneficial fitness flux is more sensitive to *U*_*b*_ than deleterious fitness flux is to *U*_*d*_, increasing the total mutation rate helps the population adapt faster. The counterintuitively increased rate of adaptation directly contradicts dysgenic fears about the consequences of elevated mutation rates on mean population fitness load.

Beneficial fitness flux might be recombination limited ^67^. Since most meioses involve exactly two crossovers per chromosome ^46^, the primary determinant of *genome-wide* recombination rate (as opposed to local recombination rates) is the number of chromosomes. Supplementary Figure 4 shows that adding more chromosomes would not substantially speed human adaptation. However, having fewer than 10 chromosomes would substantially slow down adaptation, independently of mutation rate. Humans are well above this threshold number of chromosomes, but a few of our primate relatives (the lemur *Lepilemur mustelinus*, and the collared titi monkey *Cheracebus torquatus*) are not ^68^.

Perhaps the most striking impact of high human *U*_*d*_∼2 − 10 is to create high variance in load among individuals. Figure 4 depicts this variance, given log-normally distributed fitness, in terms of the fold-difference in fitness between two individuals that are one standard deviation apart on a log scale. This difference of 15%-40% (in the historical human environment) is relatively insensitive to *s*_*b*_ and *U*_*b*_, but depends dramatically on *U*_*d*_. Adaptation time is inversely proportional to census *N*, making its sensitivity to *U*_*d*_ independent of *N* (Supplementary Table 1). The high variance among individuals engendered by high *U*_*d*_ is nearly independent of *N* (Supplementary Table 2). High within-population variance can alternatively be expressed as high mutation load 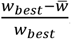.

**Figure 4.**
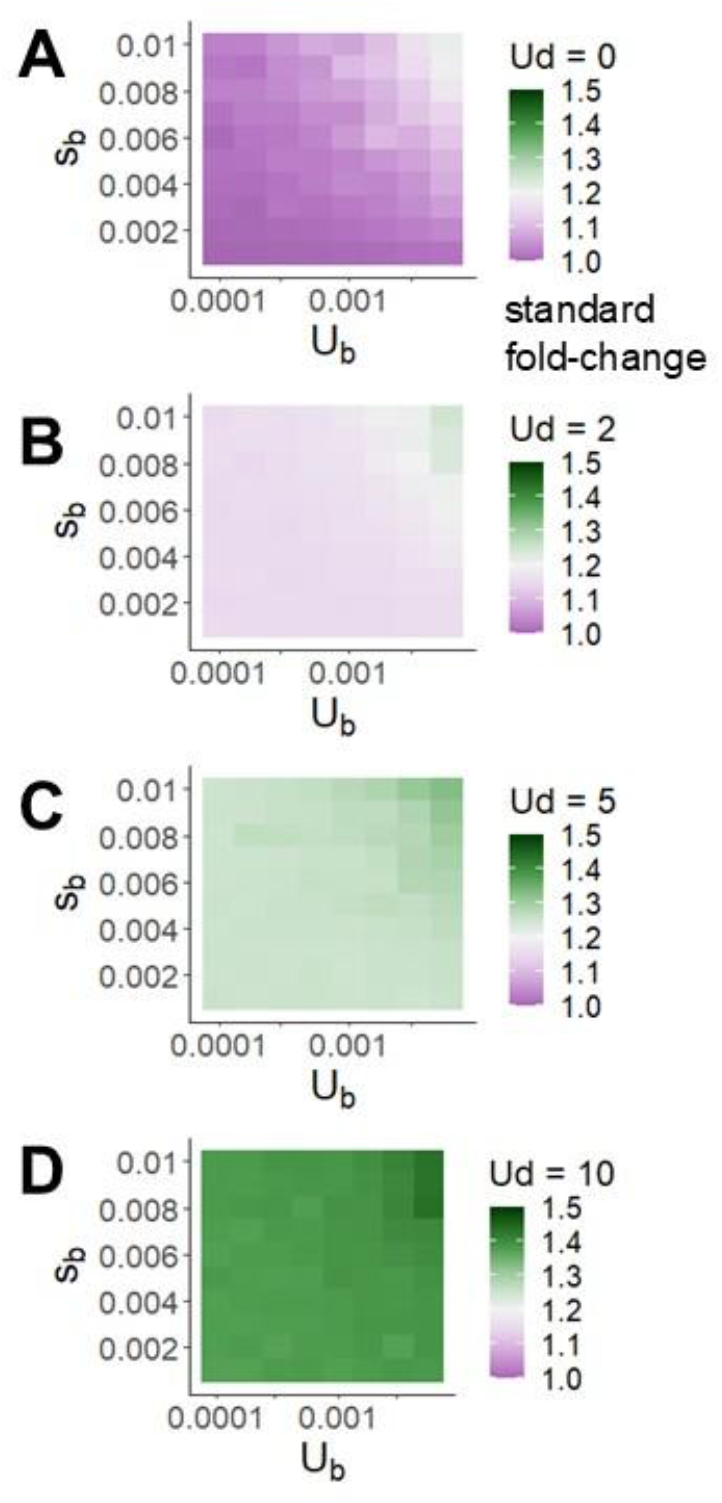
Higher deleterious mutation rates produce much more within-population variation in fitness, shown here as the fold-change in fitness corresponding to one standard deviation in the natural logarithm of fitness. The highest beneficial mutation rates and beneficial effect sizes we consider also affect within-population variation, but to a much smaller degree.

Figure 5 shows analytical expectations of high load, in the absence of linkage disequilibrium, when empirically inferred values of 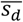 (red shading ^55,69^) as well as *U*_*d*_ are used. Grey shading shows how the unrealistically low values of 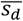 used by Galeota-Sprung *et al*. ^26^ caused them to underestimate load. In our simulations, linkage disequilibrium drives variance in fitness modestly higher again than analytical expectations of 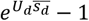 from Galeota-Sprung *et al*. ^26^ (Supplementary Figure 5). Very high values of both *U*_*b*_ and *s*_*b*_ are needed for a substantial further increase in fitness variance (Supplementary Figure 5).

**Figure 5.**
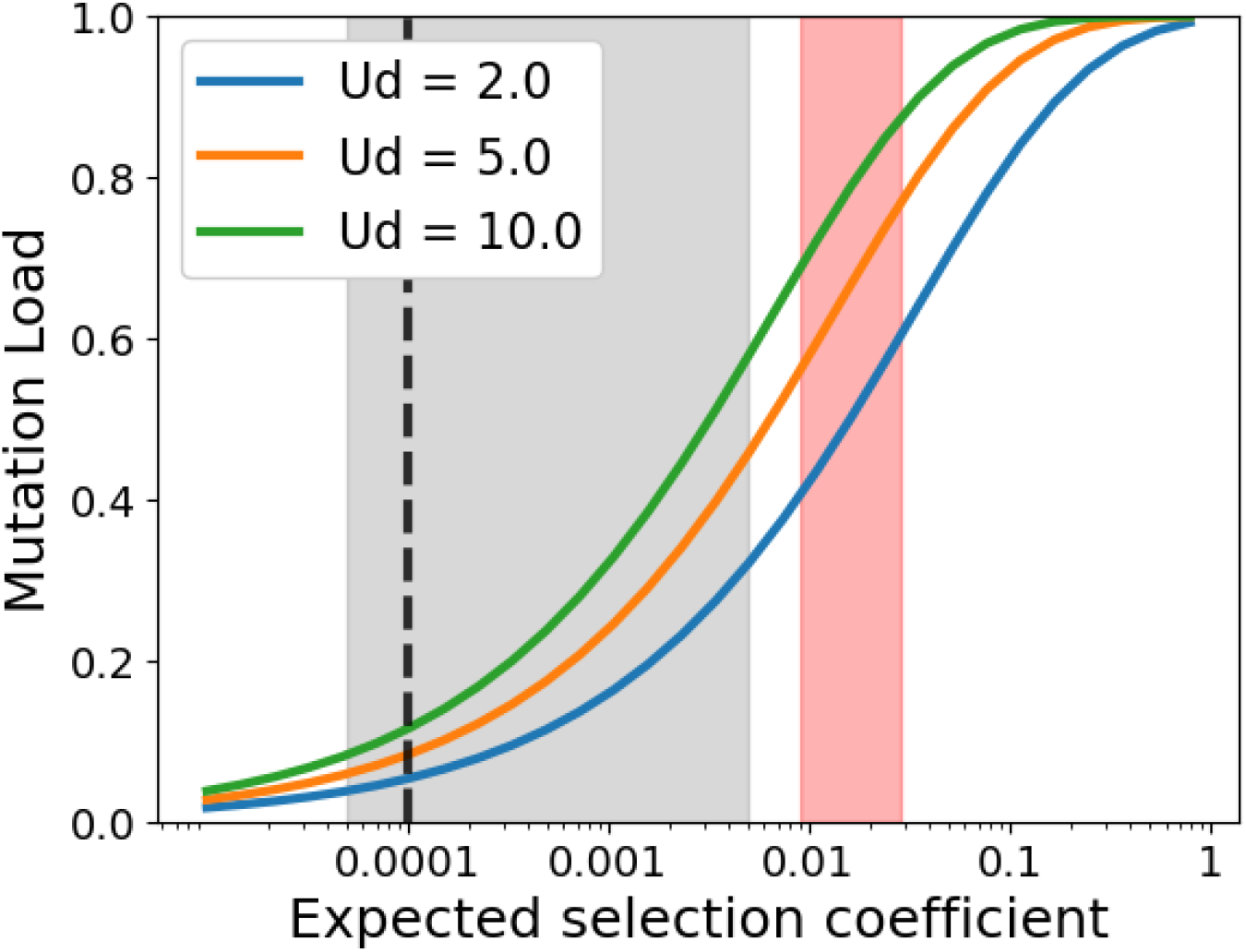
Mutation load 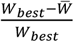 in humans is high in the absence of beneficial mutations, linkage disequilibrium, or epistasis. The mean and variance in the number of deleterious polymorphisms per individual are both *U*_*d*_/*s*_*d*_ ^70,71^, and the resulting variance in fitness is 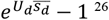. When mean population fitness is 1, the best individual present then has fitness 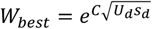.We approximate *C ≈ μ*_*G*_ + *γσ*_*G*_ from a Gumbel distribution with location 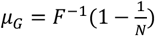 and scale parameter 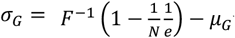 where *F*^−1^ is the normal distribution’s quantile function, *N* is population size, *γ* is the Euler-Mascheroni constant and *e* is Euler’s number ^26^. We use *N* = 10,000. Red shaded area corresponds to 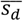 (or more strictly speaking, the heterozygotic effect size 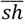) between 0.009 and 0.029, as inferred for humans by Kim *et al*. ^55^ and Boyko *et al*. ^69^, respectively. Gray shaded area corresponds to values considered by Galeota-Sprung *et al*. ^26^. Charlesworth ^72^ assumed *sN*∼1, black dashed line, corresponding to similarly low *s*∼0.0001 for humans.

## Discussion

The accumulation of many slightly deleterious fixations, driven by a realistically high human genome-wide deleterious mutation rate *U*_*d*_ = 2 − 10, is counteracted asymmetrically by a smaller number of beneficial fixations of larger effect size. We simulate this balance in the presence of complex linkage disequilibria, given conservatively low beneficial parameter values, combined with known human values for recombination and deleterious effect sizes. While population persistence is achieved, large within-population variance in fitness is created. Much of the resulting adaptation (as per Fisher’s Fundamental Theorem ^73^) is absorbed by counteracting deleterious fixations, and so despite high fitness variance, adaptation to a changing environment is 13-55% slower.

Understanding how mutation load is stabilized in humans and similar species is a precondition for addressing a long-standing fear: that recent changes to human lifestyles or technology will push us to higher load. For example, if the human mutation rate, beginning at an already critically high level, were to increase due to increased paternal age, or if selection against deleterious mutations were relaxed due to modern medicine, the perception has been that population mean fitness would decrease, potentially with disastrous consequences ^2,3,6^. Intriguingly, our results suggest that the expected increase in mutation rates in human populations due to increased paternal age has the opposite effect, improving rather than degrading population mean fitness.

High *U*_*d*_ does not threaten population persistence, but does have profound implications for within-population variation among individuals. While a genotype whose load used to cause a ∼30% reduction in fitness in ancient human environments might now have a lesser impact on fitness, it likely still has a significant impact on health. Indeed, variation in self-reported health has a substantial genetic component ^74^, and load, as assessable from whole-genome sequencing, can be used to predict medically relevant phenotypes ^4,75^. High genetic variance among individuals is a hidden confounding variable in a vast range of studies ^76^, including many studies of human health. Our theoretical assessment, when combined with estimates of human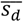, implies high variance in human mutation load, requiring a significant reassessment across all public health studies grounded in correlational analysis ^76^. A remaining caveat here is our neglect of epistasis. Positive epistasis is expected to increase variance and negative epistasis to decrease it ^77^.

High fitness variance (squared differences in fitness) is closely related to historical debates about high mutation load (differences in fitness). Early discussions of high mutation load have been critiqued as comparing mean fitness to that of a non-existing ideal genotype ^78^, one that represents invalid typological thinking ^79^. Correctly defining load as a comparison between mean population fitness and the best genotype present avoids this criticism and necessarily reduces the estimated load, which has been presented as a ‘solution’ to the mutation load ‘problem’ ^26,72,78^. While Galeota-Sprung et al. ^26^ argued that loads are small when thus conceived, their argument (see their Table 1) requires smaller 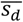 than that inferred for humans (Figure 5).

Our simulations confirm that the solution to population persistence is a pattern of many small mutations, each of which cannot be effectively cleared, being counteracted by compensatory mutations with global effects. This pattern is part of drift barrier theory ^23,24,80–87^. Drift barrier theory (Figure 6; blue circle) emphasizes the causal importance of census population size in producing a ratchet of increasing molecular and organismal complexity. Lynch ^84^ (see Chapter 4) posits that the minimum size of a deleterious mutation that can be reliably purged (‘effective population size’) is driven (albeit not exclusively) by census population size, which is in turn driven by life history traits such as body size. A low effective population size that cannot purge small DNA insertions leads to a bloated genome, whose complexity is posited to lead to larger body size and/or increased ecological specialization, reducing census population size, closing the causal loop. Drift barrier theory sees increased mutation rate as a consequence of relaxed selection against mutator alleles.

**Figure 6.**
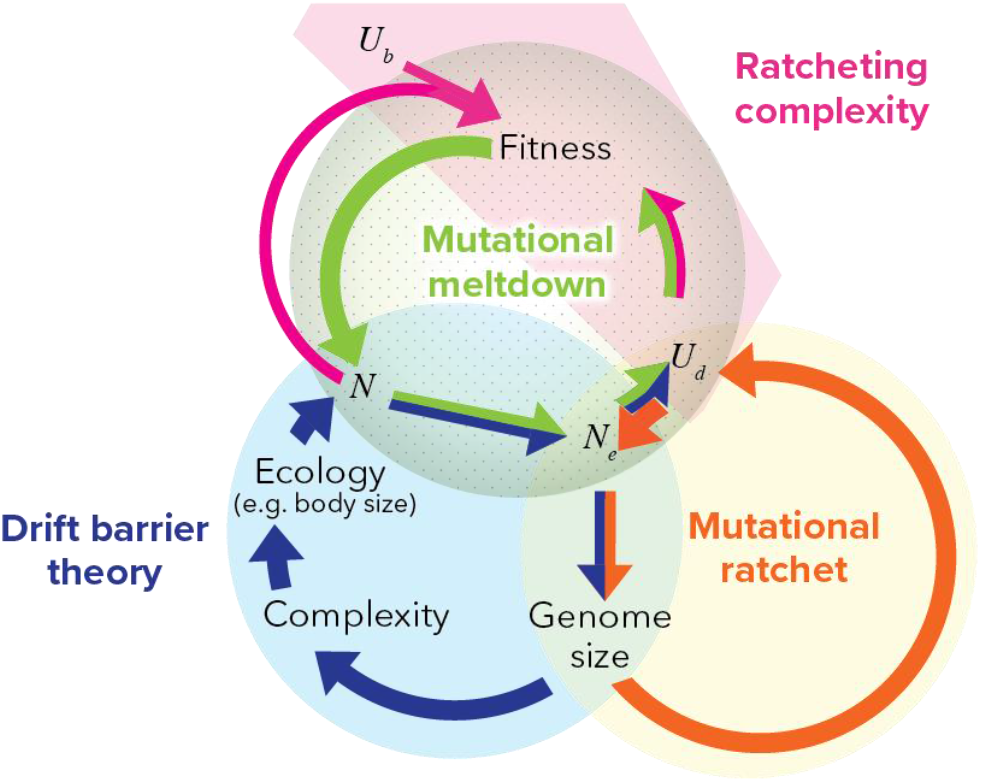
A feedback loop of ratcheting complexity can be driven either by census size *N* and ecology (drift barrier theory, blue) or by high deleterious mutation rate *U*_*d*_ (our view, orange and pink). The drift barrier ratchet requires low census population size *N*, whereas our ratchet requires high deleterious mutation rate *U*_*d*_. Drift barrier theory emphasizes a causal link from *N*_*e*_ to *U*_*d*_ via relaxed selection against mutators ^84^, whereas we emphasize background selection as a causal driver in the opposite direction, i.e. from *U*_*d*_ to *N*_*e*_ . Mutational meltdown ^93,94^ is shown for completeness (green), since its elements are already invoked.

We suggest shifting the causal emphasis to high deleterious mutation rate instead of low census population size. Indeed, a mutational ratchet (Figure 6, orange cycle) can occur even when census population size is high. First, high *U*_*d*_ > 1 substantially reduces *N*_*e*_ through background selection (among unlinked sites) ^28,88^ and via linkage disequilibrium with beneficial mutations ^29^. As with drift barrier theory, the resulting deluge of slightly deleterious fixations increases genome size ^89^ — note that a ratchet will not occur if deleterious fixations decrease genome size, as is likely the case in prokaryotes ^90–92^. The feedback loop from there does not go through census *N*; instead, larger genomes create a larger target size for deleterious mutations, directly increasing *U*_*d*_.

The mutational ratchet described above (Figure 6, orange) might drive *U*_*d*_ up to a high enough level to power the complexity ratchet (Figure 6, pink) that is the focus of this manuscript. Similarly to drift barrier theory, when slightly deleterious fixations cannot be reversed to achieve detailed balance, they are compensated for by large effect changes that frequently occur at a higher level of organization, creating a complexity ratchet. The key difference is that the drift barrier theory (Figure 6, blue and pink) requires low *N* but can occur at low *U*_*d*_, so long as *sN*_*e*_ is low. Our alternative (Figure 6, orange and pink) requires *U*_*d*_ > 1, and can occur even for high census *N*.

Previous hypotheses have focused on population size as the crucial difference between species that are able to purge load within a small, streamlined genome (e.g. bacteria) vs. species forced into ratcheting molecular complexity in search of innovative molecular solutions to stay ahead of perpetual degradation (e.g. humans). But differences in *U*_*d*_ could also explain differences in molecular complexity across the tree of life.

We confirmed that *U*_*d*_ > 1 need not cause population decline, because the accumulation of mildly deleterious fixations (Ohta’s ratchet) is countered by a smaller number of larger-effect beneficial fixations under quite conservative assumptions. While *U*_*d*_ > 1 may therefore not threaten population persistence, it may be an important driver of the evolution of molecular and organismal complexity. Combined with realistic deleterious effect sizes, high *U*_*d*_ created high fitness variance within human populations in their ancestral environment.

## Code Availability

Simulation code in C is available at github.com/MaselLab/MutationLoad.

## Acknowledgements

We thank Dr. Kun Xiong for helping the first author learn C, Dr. Walid Mawass for helpful discussions, and Dr. Cristian Román-Palacios for assistance navigating the database of chromosome numbers across the tree of life. This work was supported by the John Templeton Foundation [62028] and NIH Grant GM084905. Figure 6 was created by Jennifer Yamnitz.

