## Supplemental File for "Human deleterious mutation rate slows adaptation and implies high fitness variance"

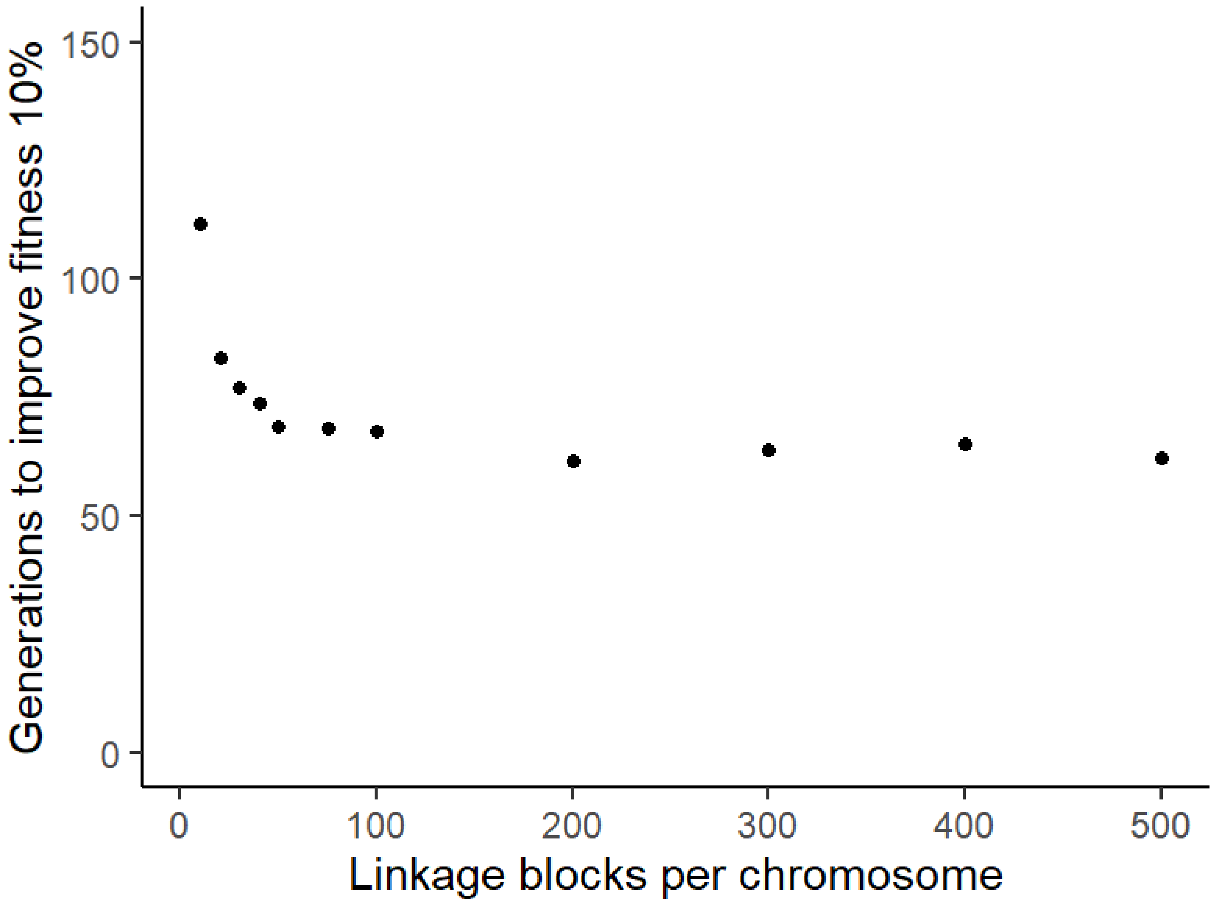


**Supplementary Figure 1.** Dividing each of our 23 chromosomes into fewer than 50 linkage blocks would substantially underestimate adaptation, but using more linkage blocks has only small benefits. All have $N = 20{,}000$, $U_d = 5$, $U_b = 0.001$, and mean $s_b = 0.005$.

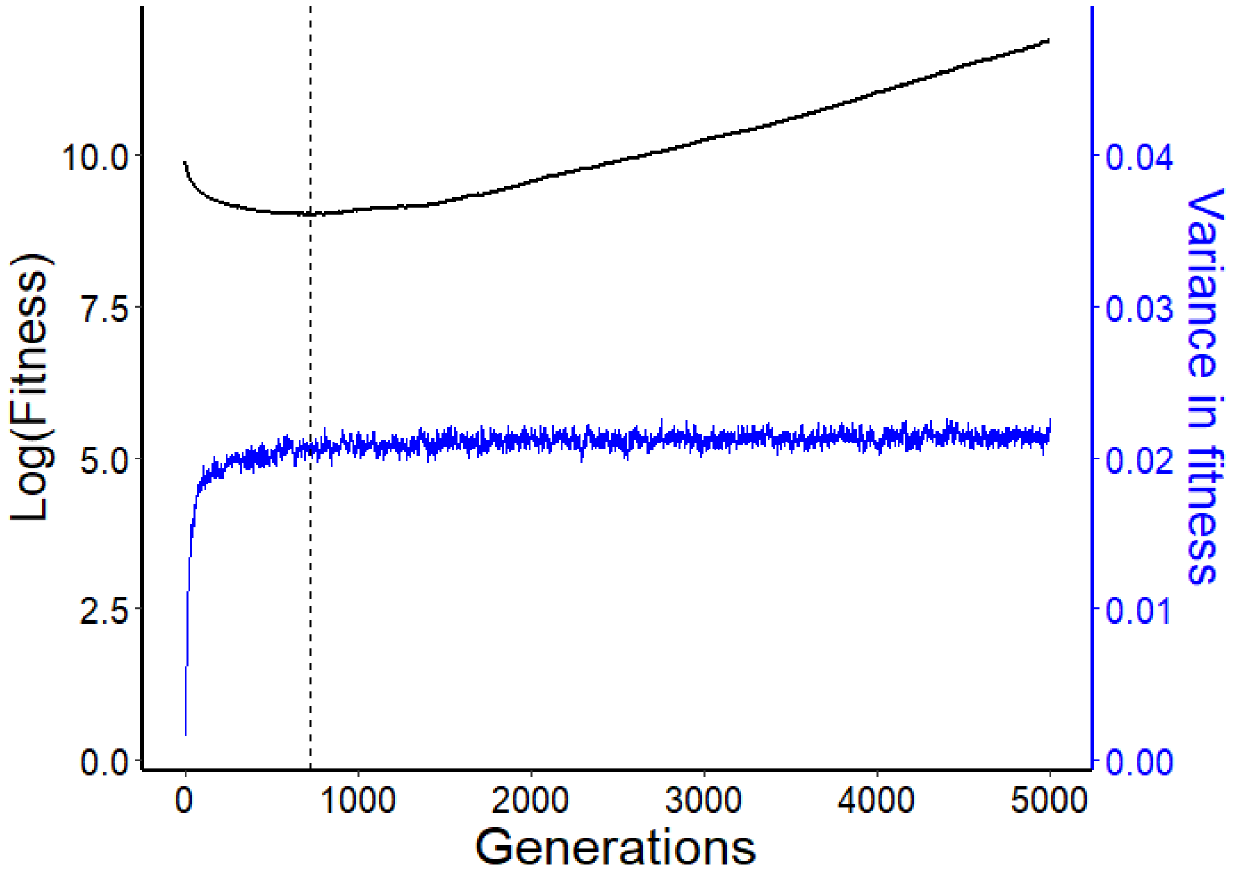


**Supplementary Figure 2. The burn-in phase ends once variance in fitness plateaus.** Dashed line indicates the end of the burn-in phase, at generation 1227. The slope of log mean fitness (black) after the burn-in phase describes the net fitness flux in the population over time. N = 20,000, each individual has 23 chromosomes with 50 linkage blocks per chromosome, $U_b$ =0.002, and $s_b$ = 0.0025.

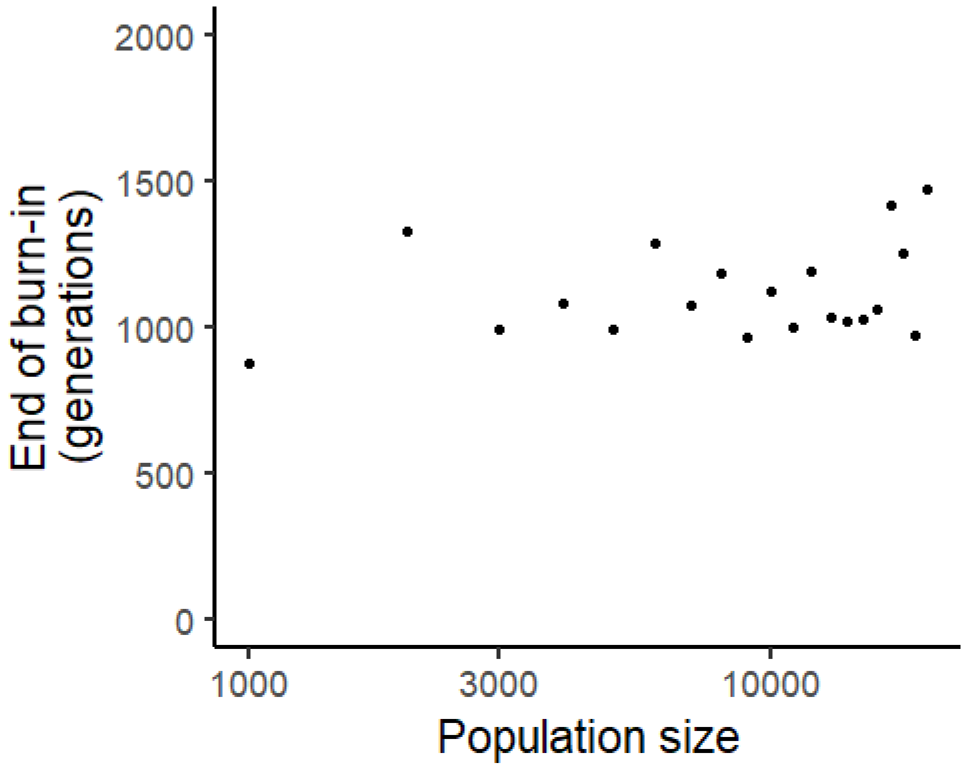


**Supplementary Figure 3.** The number of generations until the end of the burn-in phase increases slowly with increasing population size.

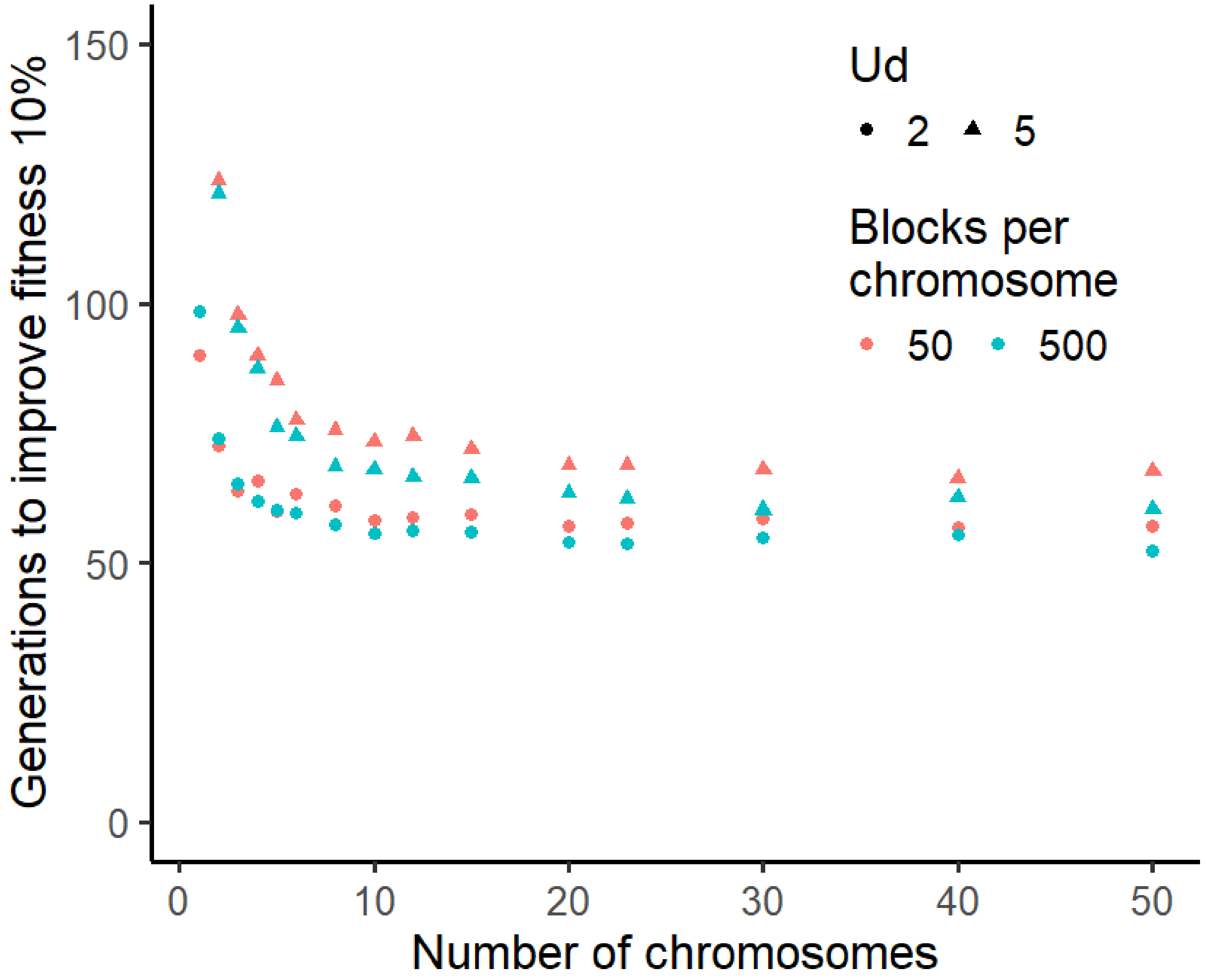


**Supplementary Figure 4.** Below a threshold number of chromosomes (~10), adaptation under a human parameter value regime is substantially slowed by lack of recombination. The location of the threshold is independent of $U_d$ within the range 2 – 10 (not shown). We used 500 linkage blocks per chromosome, rather than the 50 used for computational convenience elsewhere, to avoid underestimating the power of recombination, especially for fewer chromosomes. $N = 20{,}000$, $U_b = 0.001$, and $\bar{s_b} = 0.005$.

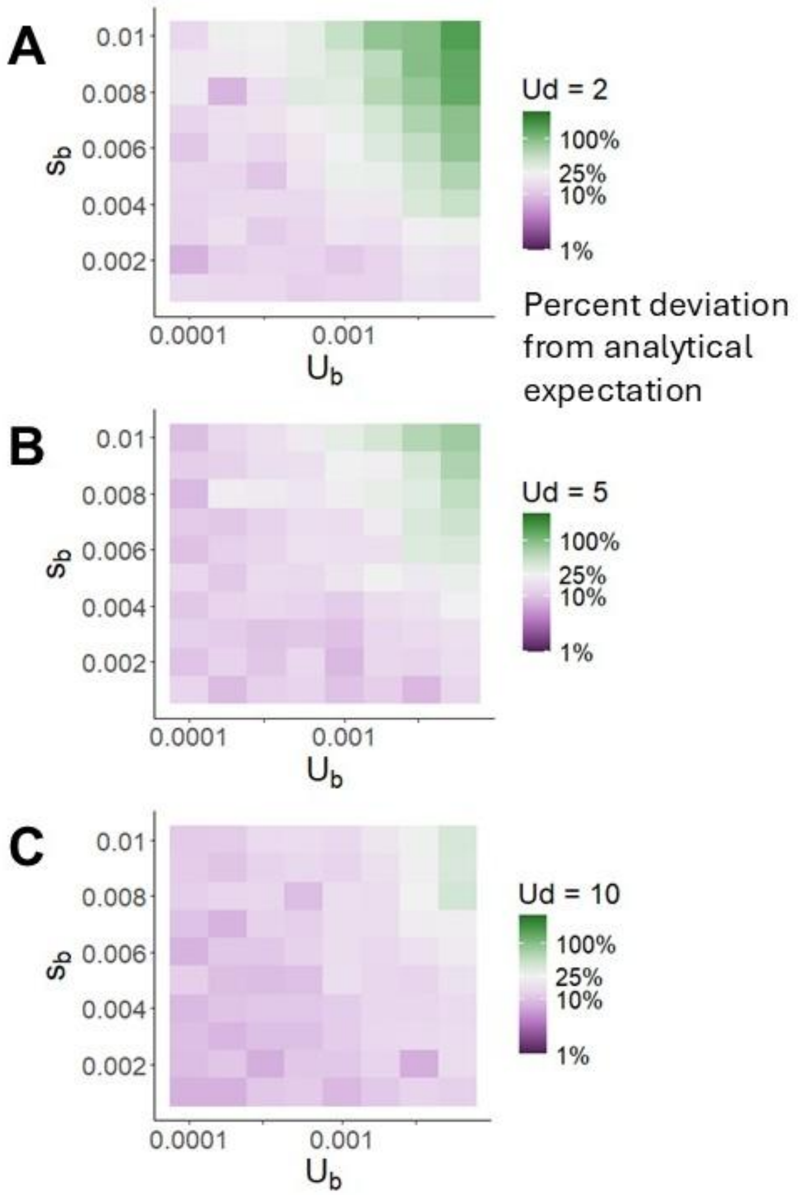


**Supplementary Figure 5.** The presence of beneficial mutations and emergent linkage disequilibrium increases variance above the analytical expectation $e^{U_d\overline{s_d}} - 1$ from Galeota-Sprung *et al*. (2020). However, the increase in variance is <50% of the analytically expected variance for all but the highest beneficial mutation rates and mean effect sizes, and <25% of the expected variance (shown as purple rather than green) for most of the beneficial parameter value range we consider. The proportional effect of beneficial mutations on fitness variance is less pronounced when baseline variance is higher at higher deleterious mutation rates.

| | $N = 10{,}000$ | $N = 20{,}000$ | $N = 40{,}000$ |
|---|---|---|---|
| $U_d = 0$ | 104.4 | 51.21 | 25.73 |
| $U_d = 2$ | 111.7 | 57.62 | 30.47 |
| $U_d = 5$ | 145.5 | 68.82 | 36.32 |
| $U_d = 10$ | 281.2 | 104.05 | 48.68 |

**Supplementary Table 1.** The number of generations of adaptation needed to improve mean relative fitness by 10% in a population with $U_b = 0.001$ and $s_b = 0.005$. Adaptation time is approximately linearly dependent on $N$, making the degree of sensitivity to $U_d$ independent of $N$.

| | $N = 10{,}000$ | $N = 20{,}000$ | $N = 40{,}000$ |
|---|---|---|---|
| $U_d = 0$ | 1.029 | 1.040 | 1.067 |
| $U_d = 2$ | 1.160 | 1.165 | 1.166 |
| $U_d = 5$ | 1.262 | 1.263 | 1.266 |
| $U_d = 10$ | 1.386 | 1.392 | 1.384 |

**Supplementary Table 2.** The standard deviation in the natural logarithm of fitness, expressed as a fold-change factor, at the end of a simulation in a population with $U_b = 0.001$ and $s_b = 0.005$. For high $U_d$, there is little dependence on $N$.
